# Alternative lipid synthesis in response to phosphate limitation promotes antibiotic tolerance in Gram-negative ESKAPE pathogens

**DOI:** 10.1101/2024.09.11.612458

**Authors:** Roberto Jhonatan Olea-Ozuna, Melanie J. Campbell, Samantha Y. Quintanilla, Sinjini Nandy, Jennifer S. Brodbelt, Joseph M. Boll

## Abstract

The Gram-negative outer membrane protects bacterial cells from environmental toxins such as antibiotics. The outer membrane lipid bilayer is asymmetric; while glycerophospholipids compose the periplasmic facing leaflet, the surface layer is enriched with phosphate-containing lipopolysaccharides. The anionic phosphates that decorate the cell surface promote electrostatic interactions with cationic antimicrobial peptides such as colistin, allowing them to penetrate the bilayer, form pores, and lyse the cell. Colistin is prescribed as a last-line therapy to treat multidrug-resistant Gram-negative infections.

*Acinetobacter baumannii* is an ESKAPE pathogen that rapidly develops resistance to antibiotics and persists for extended periods in the host or on abiotic surfaces. Survival in environmental stress such as phosphate scarcity, represents a clinically significant challenge for nosocomial pathogens. In the face of phosphate starvation, certain bacteria encode adaptive strategies, including the substitution of glycerophospholipids with phosphorus-free lipids. In bacteria, phosphatidylethanolamine, phosphatidylglycerol, and cardiolipin are conserved glycerophospholipids that can form lipid bilayers, particularly in the presence of other lipids. Here, we demonstrate that in response to phosphate limitation, conserved regulatory mechanisms induce alternative lipid production in *A. baumannii*. Specifically, phosphate limitation induces formation of three lipids, including amine-containing ornithine and lysine aminolipids. Mutations that inactivate aminolipid biosynthesis exhibit fitness defects relative to wild type in colistin growth and killing assays. Furthermore, we show that other Gram-negative ESKAPE pathogens accumulate aminolipids under phosphate limiting growth conditions, suggesting aminolipid biosynthesis may represent a broad strategy to overcome cationic antimicrobial peptide-mediated killing.

**Author Summary:** Gram-negative ESKAPE pathogens, including A*cinetobacter baumannii*, are responsible for a dramatic increase in the morbidity and mortality of patients in healthcare settings over the past two decades. Infections are difficult to treat due to antibiotic resistance and tolerance; however, broadly conserved mechanisms that promote antibiotic treatment failure have not been extensively studied. Herein, we identify an alternative lipid biosynthesis pathway that is induced in phosphate starvation that enables Gram-negative ESKAPE pathogens, including *A. baumannii*, *Klebsiella pneumoniae*, and *Enterobacter cloacae* to build lipid bilayers in the absence of glycerophospholipids, which are the canonical bilayers lipid. Replacement of the anionic phosphate in the lipid headgroup with zwitterionic ornithine and lysine promote survival against colistin, a last resort antimicrobial used against Gram-negative infections. These studies suggest that ESKAPE pathogens can remodel their bilayers with phosphate free lipids to overcome colistin treatment and that aminolipid biosynthesis could be targeted to improve antimicrobial treatment.

## Introduction

The Gram-negative cell envelope consists of a symmetrical bilayer of glycerophospholipids in the inner membrane, while the outer membrane exhibits an asymmetrical composition, with glycerophospholipids in the periplasmic leaflet and lipopolysaccharide enriched in the outer leaflet (1). The intricate organization underscores the remarkable complexity of bacterial membrane architecture, crucial for microbial survival in various environments. However, under specific stress conditions such as nutrient limitation, temperature fluctuations or exposure to antimicrobial agents, certain bacteria activate alternative lipid biosynthesis pathways or modify existing lipids to adapt and ensure cellular viability (2). One example is aminolipids, which contain amino acid headgroups like lysine, glycine, glutamine, and serine-glycine, with ornithine being the most common (3,4). Ornithine lipids (OLs) are phosphorus-free and found exclusively in bacteria; they are absent in archaea or eukaryotes (5). Their basic structure comprises a 3-hydroxylated fatty acid linked by an amide bond to the α-amino group of ornithine and a second fatty acid attached by an ester bond to the 3-hydroxyl group of the first fatty acid (6). Although OLs are found in both the inner and outer lipid bilayers of Gram-negative bacteria, they are enriched in the outer membrane (7–10). OL biosynthesis is catalyzed by two acyltransferases, OlsB and OlsA, or by the bifunctional acyltransferase OlsF (11–13). In some Gram-negative pathogens such as *Pseudomonas aeruginosa* or *Vibrio cholerae*, OLs are exclusively formed under phosphate limiting conditions (14,15), indicating the presence of a specific regulatory mechanism. The importance of aminolipids transcends basic physiology, especially in the context of antibiotic resistance. ESKAPE pathogens, a group of pathogens that include *Acinetobacter baumannii*, are notorious for their ability to overcome antibiotic treatment and cause hospital-acquired infections (16). Aminolipid synthesis has been implicated in increased bacterial fitness under antimicrobial stress (9,17,18), potentially contributing to pathogen persistence in clinical settings. Additionally, there is a notable relationship between membrane lipid remodeling and resistance to colistin, a last-resort antibiotic that is used against multi-drug resistant Gram-negative infections (19–22). Chemical modifications to the lipid A domain of lipopolysaccharide or enrichment of amino acid-containing glycerophospholipids have been associated with colistin resistance (23–25), highlighting the importance of understanding lipid metabolism in combating antibiotic resistance.

In this study, we demonstrate that *A. baumannii* produced two aminolipids in limiting phosphate growth conditions, including lysine lipids (LLs) and OLs. OL and LL synthesis is dependent on the *olsB* and *olsA* genes, and *olsB* expression is regulated transcriptionally by the response regulator, PhoR. Additionally, mutants deficient in aminolipid synthesis exhibit increased colistin susceptibility relative to wild type. We also found that other Gram-negative ESKAPE pathogens, including *Klebsiella pneumoniae* and *Enterobacter cloacae*, accumulate aminolipids under phosphate limited growth conditions. These findings suggest a broad survival strategy among ESKAPE pathogens that could promote survival during antibiotic treatment.

## Results

### Phosphate limitation induces lipid membrane composition modifications in *A. baumannii*

The membrane lipid composition of diverse *A. baumannii* isolates was analyzed, including strains ATCC 17978, ATCC 19606, and AB5075, cultivated in complex lysogeny broth (LB) medium supplemented with ^32^P-orthophosphoric acid. Labelled cells were collected at mid-logarithmic growth phase, lipids were extracted using the Bligh and Dyer method, and separated by hydrophobicity using two-dimensional thin-layer chromatography (TLC), as previously (20,26,27). We also prepared lipid extracts from *Escherichia coli* K-12 strain W3110 for comparison, which served as a well-characterized standard (28). TLC analysis showed conserved glycerophospholipid enrichment corresponding to known structures, including phosphatidylethanolamine (PE), phosphatidylglycerol (PG), and cardiolipin (CL) (**Figure S1A**). Additionally, *A. baumannii* strains also produced two distinct lipid species, including lyso-PE (20) and an unknown phospholipid, denoted as UPL1, that could be a CL derivative, mono-lyso CL (29). The chemical structures of known phospholipids are shown in **Figure S1B**.

*A. baumannii* strain ATCC 17978 was cultured in minimal medium with either 1 mM (excess) or 50 μM (limiting) phosphate, conditions commonly used to distinguish between phosphate-sufficient and phosphate-limited growth (6, 13–15, 29, 30). Growth in 1 mM phosphate induced a lipid composition almost equivalent to growth in complex LB medium (**Figure 1 and Figure S1A**). One notable change was that UPL1 was absent, and another unidentified lipid, denoted as UPL2, was formed. To explore lipid biosynthesis under phosphate-limiting conditions, membrane lipid profiles were analyzed after growth in minimal media supplemented with 50 μM (limiting) phosphate concentrations (**Figure 1**). Limited phosphate availability impacted relative lipid levels, suggesting decreased phosphate-containing lipid synthesis. However, unlike PE and PG glycerophospholipids, relative CL levels did not change significantly, suggesting its production may provide a strategy to maintain membrane charge stability. Concomitant production of three unknown lipids, referred to as unknown lipids 1 (U1), 2 (U2), and 3 (U3), were produced in phosphate limiting growth. Ninhydrin staining revealed that U1 and U2 contained free amines, like PE. Increased ratios of ninhydrin-stained unknown aminolipids relative to PE showed that under phosphate-limiting conditions, potential phosphate-free aminolipids were produced. The total lipid quantifications after lipids were extracted and separated on TLC is shown in **Figure S2A.**

**Figure 1:**
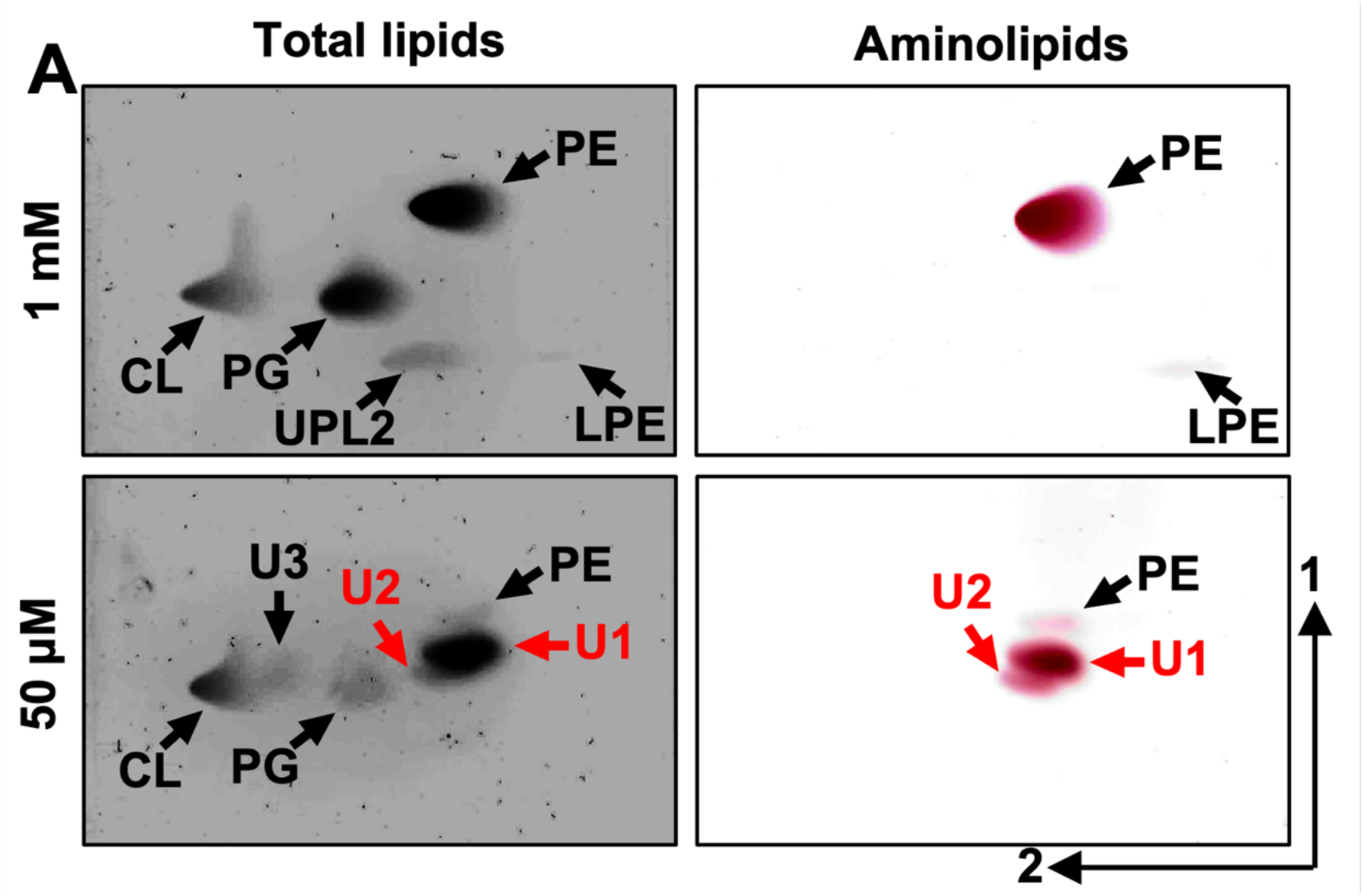
Lipid composition of *A. baumannii* strain ATCC 17978 in excess and limiting phosphate concentrations. **A.** Strains were grown in basal media 2 (BM2) with excess (1mM) or limiting (50 µM) phosphate. Cells were collected, lipids were extracted using the Bligh and Dyer method and separated using 2-dimensional thin-layer chromatography. Total lipids were stained with sulfuric acid (left). Aminolipids were stained using ninhydrin (right). Specific lipids are labelled: PE, phosphatidylethanolamine; PG, phosphatidylglycerol; CL, cardiolipin; LPE, lyso-PE; U1, unknown lipid 1; U2, unknown lipid 2; UPL1, unknown phospholipid 1. Red letters denote aminolipids that provide the focus of the study.

Growth in limiting phosphate concentrations slowed growth in ATCC 17978 relative to excess phosphate (**Figure S2B**), and microscopic analysis revealed that cells elongated and increased their surface area when phosphate is limiting (**Figure S2C and S2D**), a response previously reported in other Gram-negative bacteria (32). Additionally, while the composition of lipooligosaccharide (LOS) fractions remained consistent across limiting phosphate conditions, the relative level of LOS was decreased under phosphate limitation (**Figure S2E**).

### Aminolipids synthesized during phosphate limitation are OLs and LLs

One dimensional TLC showed three lipids (PE, U1, and U2) stained with ninhydrin (**Figure S3A**). After separation of lipids based on hydrophobicity, individual bands corresponding with U1 and U2 were scraped from the TLC plates and extracted using the Bligh and Dyer method. U1 and U2 bands were analyzed by liquid chromatography-mass spectrometry (LC-MS) and structurally characterized using tandem mass spectrometry (MS/MS) (**Figure 2**). The elution profiles obtained for the U1 and U2 bands are displayed in **Figures 2A** and **2D**, respectively. To determine the composition of aminolipids, a data-dependent acquisition method was used to isolate and activate the most abundant ions detected in each chromatographic peak with higher energy collisional dissociation (HCD) in negative-ionization mode. HCD of precursor *m/z* 621.52 found in the U1 extract resulted in the loss of the headgroup, which was observed at *m/z* 131.08, and corresponded to deprotonated ornithine (**Figure 2B**). The most abundant fragment ion (*m*/*z* 367.30) was used to identify the acyl chain connected to the headgroup as 16:0 (number of carbon atoms: double bonds). Further, the complementary ion of *m*/*z* 253.22 confirmed the identity of the fatty acid connected at the 3-hydroxyl position of the first fatty acid as 16:1. Following this analysis, lipids containing a double bond were subsequently targeted in a second LC run in the positive-ionization mode using 193 nm ultraviolet photodissociation (UVPD) for MS/MS. UVPD is an alternative fragmentation method that utilizes high-energy photons to activate and dissociate the selected lipid precursor ions, allowing localization of the double bonds within the fatty acyl chains. UVPD of precursor *m/z* 623.53 produced two fragment ions separated by 24 Da that originate from cleavages adjacent to the double bond. This pair of diagnostic ions localizes the double bond to the 9^th^ position (**Figure 2C**). This LC-MS/MS strategy identified 49 OLs, including double bond isomers, and 10 unknown lipids in the U1 extract (**Table S1**) and a total of 24 OLs and 16 unknown lipids in the U2 extract (**Table S2**). HCD of the unknown lipids yielded similar fragmentation to that observed for OLs, but the fragmentation patterns were distinguished by the release of a deprotonated headgroup that corresponded to either a lysine or monomethylated ornithine headgroup (*m*/*z* 145.10) (**Figure 2E**). While the aminolipid head group could be either lysine or monomethylated ornithine, the absence of an ortholog for OlsG, the enzyme responsible for OL methylation (33), and the presence of *olsG* only in certain planctomycete genomes, strongly suggests that OL methylation is unlikely to occur in *A. baumannii*. Therefore, the identified lipid is denoted herein as a LL.

**Figure 2:**
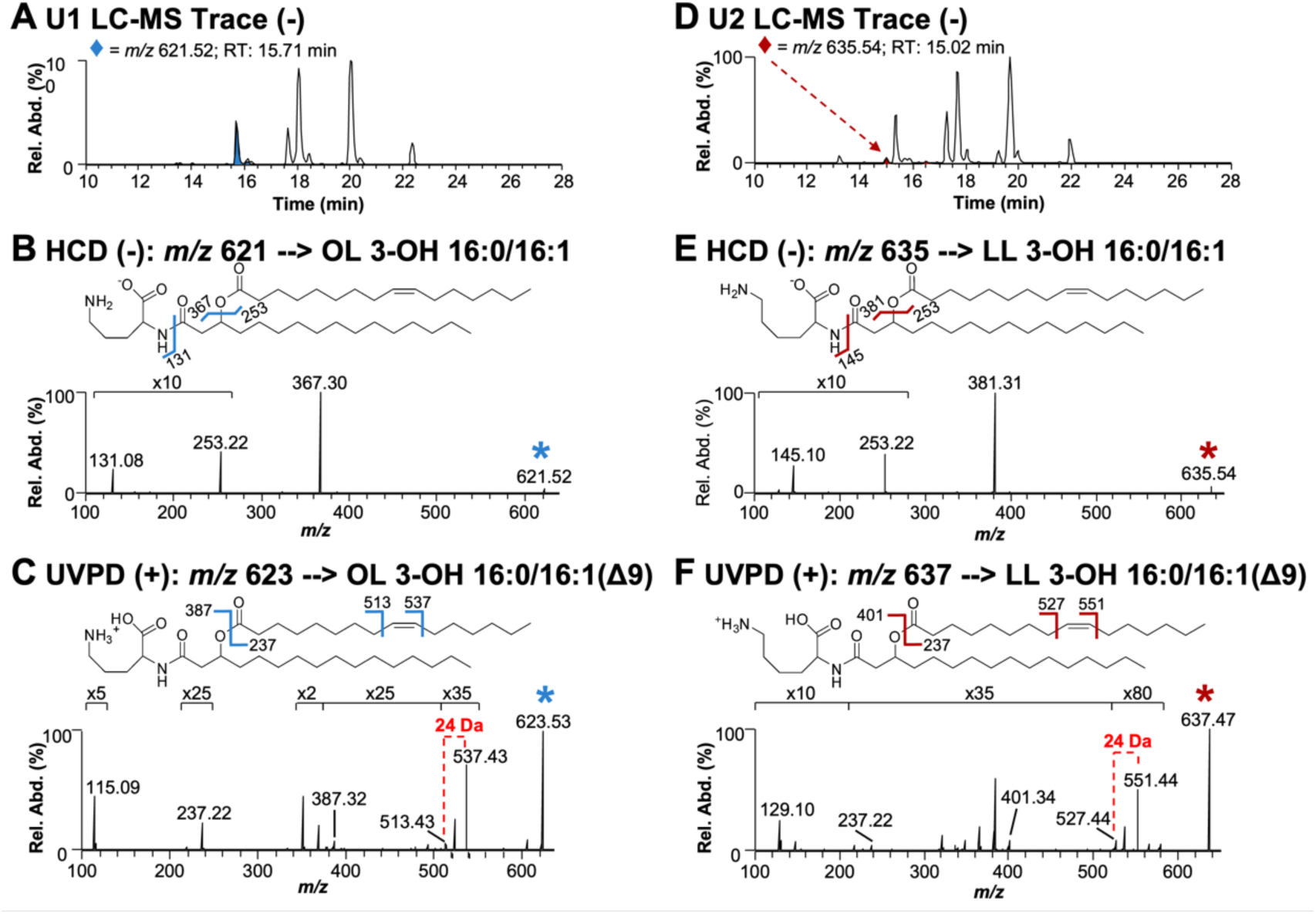
Structural analysis of the lipids produced during phosphate limitation. **A.** LC-MS trace of U1 lipid extract in negative-ionization mode. **B.** HCD (NCE 22) mass spectrum of *m/z* 621.52 ([M-H]^-^), an ornithine lipid found in the U1 extract**. C.** UVPD (4 pulses at 2 mJ/pulse) mass spectrum of *m/z* 623.53 ([M+H]^+^). **D.** LC-MS trace of U2 lipid extract in negative-ionization mode. **E.** HCD (NCE 22) mass spectrum of *m/z* 635.54 ([M-H]^-^), a lysine lipid found in U2 extract. **F.** UVPD (4 pulses at 2 mJ/pulse) mass spectrum of *m/z* 637.47 ([M+H]^+^). The selected precursor ions are labeled with asterisks in B, C, E, and F.

After confirming that *A. baumannii* strain ATCC 17978 produces OLs and LLs, we explored if other *A. baumannii* isolates form these aminolipids during phosphate limitation. One-dimensional TLC stained with ninhydrin revealed that diverse *A. baumannii* strains, including ATCC 17978, ATCC 19606, AB5075, AYE, and the environmental isolate, *A. baylyi*, were also capable of aminolipid biosynthesis in response to phosphate limitation (**Figure S3B**). Together, these studies suggest that *Acinetobacter* can form lipid bilayers with not only glycerophospholipids, but also membranes enriched with OLs and LLs when phosphate availability is limited.

### Differentially regulated genes during phosphate starvation

To determine genes that regulate aminolipid biosynthesis, total RNA was isolated, rRNA was depleted, and transcripts were sequenced from *A. baumannii* strain ATCC 17978 after growth in minimal media supplemented with 1 mM (excess) and 50 µM (limiting) phosphate. Using a 3-fold cutoff for the weighted proportions fold change and an FDR p-value correction < 0.05, 67 upregulated genes and 109 downregulated genes were found (**Figure 3A**). Many down-regulated genes were involved in iron uptake, such as those coding for ferric uptake regulators (Fur) and components of the ABC transporters for iron. Additionally, genes involved in lysine degradation and aminoacid metabolism were down-regulated, indicating potential changes in nitrogen metabolism under phosphate-limited conditions. Upregulated genes were associated with TAT-dependent proteins export, which could be involved in the transport of key lipids or membrane proteins, as well as phospholipases, which may play a role in membrane remodeling. Furthermore, genes related to regulation and phosphate transporters were induced, likely in response to changes in phosphate availability. A complete list of significant (*P* < 0.05) up- and down-regulated genes is included in **Table S3**. The RNA-sequencing results were validated using qRT-PCR analysis to measure differential expression between three specific genes in excess and limiting phosphate concentrations (**Figure S4A**) and via phenotypic characterization of several mutants described below. Validation of the dataset suggested that phosphate limitation induces a broad transcriptional reprogramming, affecting processes such as lipid biosynthesis, membrane integrity, and nutrient uptake, which are crucial for bacterial survival under stress conditions.

**Figure 3:**
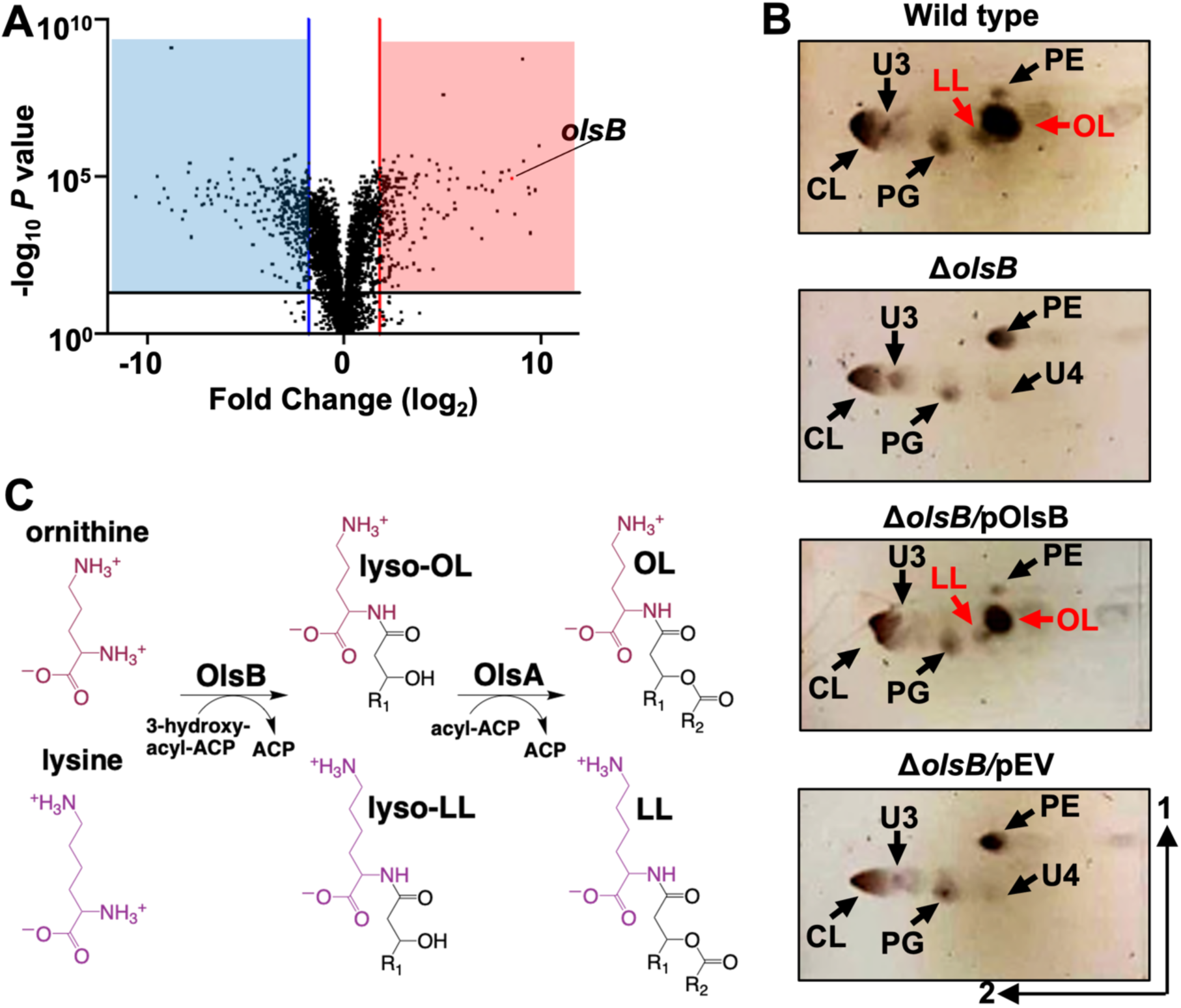
The *olsB* gene is required for ornithine and lysine lipid formation in phosphate limited growth conditions. **A.** Volcano plot of differentially regulated genes in excess and limiting phosphate. Lines indicate 3.5-fold cutoffs. **B.** 2D thin-layer chromatography stained with sulfuric acid showing wild type and Δ*olsB* strains grown in phosphate limiting (50 µM) conditions. Specific lipids are labelled: PE, phosphatidylethanolamine; PG, phosphatidylglycerol; CL, cardiolipin; OL, ornithine lipid; LL, lysine lipid; U3, unknown lipid 3; U4, unknown lipid 4. OL and LL aminolipids are labelled in red. **C.** Proposed ornithine lipid (OL) and lysine lipid (LL) biosynthesis pathways in *A. baumannii*.

One notable up-regulated gene (*A1S_0889*) showed a log_2_ fold change of 8.57 and 31% identity and 51% similarity (91% coverage) to *P. aeruginosa olsB*. While not induced in phosphate starvation, *A. baumannii* also encodes a putative *olsA* orthologue (*A1S_2990*) with 43% identity and 61% similarity (78% coverage) to *P. aeruginosa olsA*. OlsBA has been characterized in several bacteria and is involved in OLs biosynthesis (8,11,12,14,34). Notably, *A. baumannii olsA* is located at a distinct site on the chromosome relative to *olsB* and is not transcriptionally regulated by phosphate concentrations.

### *olsB* and *olsA* genes are required for aminolipids synthesis in *A. baumannii*

Differential gene expression analysis indicated changes in gene dosage in response to excess/limiting phosphate concentration. Specifically, *A1S_0889* expression increased in response to phosphate limitation, whereas *A1S_2990* expression did not change. To confirm that *A1S_0889* was an *olsB* orthologue, we generated Δ*olsB* in *A. baumannii* strain ATCC 17978, by deleting codons 34 and 214. TLC analysis revealed that under phosphate limitation, aminolipids were absent in the *olsB*-deficient mutant and a mutant carrying only the empty vector but wild-type accumulated OLs and LLs (**Figure 3B**). Furthermore, *olsB* complementation from a non-native promoter restored OL and LL formation. Based on the lipid migration patterns in wild type, the lipids are OLs and LLs.

Additionally, we generated Δ*olsB* (*HMPREF0010_01383*) using recombineering in strain ATCC 19606 and also analyzed the *olsB::tn* and *olsA::tn* mutants from the AB5075 transposon mutant library (35), which contain insertions in the respective genes, *ABUW_3039* and *ABUW_0502*. Lipid analysis when the mutants were cultivated under phosphate limitation showed that the wild-type strains ATCC 19606 and AB5075 produced OLs and LLs, while the *olsB* and *olsA* mutants did not (**Figure S4B and S4C**). The *olsA*::*tn* mutant lipid profile did not show the expected accumulation of lyso-aminolipids, consistent with findings in other Gram-negative Δ*olsA* mutants (30). This suggests that lyso-aminolipids are tightly regulated and rapidly degraded within the cell. These data suggested that ornithine and lysine lipid biosynthesis in *A. baumannii* is dependent on OlsB and OlsA. We will also note that we observed an unknown lipid, labelled as U4 in figure 3B and figure S4B that may indicate an OlsA-dependent lipid.

Together, the data supports a model suggesting that aminolipid synthesis occurs in at least two distinct steps within *A. baumannii* (**Figure 3C**), consistent with previous work in other Gram-negative bacteria (11,12). Initially, ornithine and lysine undergo acylation in an OlsB-dependent reaction, leading to the formation of lyso-OL and lyso-LL. Subsequently, in a second step facilitated by OlsA, lyso-OL and lyso-LL are further acylated at the hydroxy position, yielding OL and LL, respectively.

Microscopic analysis and comparison of LOS fractions and levels between aminolipid-producing and aminolipid-deficient *A. baumannii* strains revealed no significant morphological differences (**Figure S4D and S4E**) or changes in LOS fractions and relative levels (**Figure S2E)**. However, it is noteworthy that the core fraction of LOS from *A. baumannii* strains ATCC 19606 differs from of the ATCC 17978 or AB5075 strains. Additionally, a faint band in the core fraction was observed in the *olsA*::*tn* transposon mutant of AB5075, which was not present in the wild type strain.

### Mutants deficient in aminolipids synthesis show differential growth rate under phosphate limitation

To assess growth rates under phosphate limitation, wild-type and *olsB* mutant strains were grown in minimal medium with limiting phosphate concentrations (**Figure S4F**). The optical density at 600 nm (OD_600_) was monitored over time. While growth of strain ATCC 17978 Δ*olsB* was not impacted, the growth rate in strains ATCC 19606 and AB5075 was reduced in the *olsB* or *olsA* T101 mutants, suggesting there are strain-dependent effects on aminolipid biosynthesis that impact fitness.

### Aminolipid biosynthesis mutants are defective in colistin tolerance

Changes in lipid composition could alter the physicochemical properties of the bilayers, particularly the charge. To explore this concept, the impact of aminolipids biosynthesis on colistin tolerance was measured in *A. baumannii* strain ATCC 17978, where growth rate was not impacted in the *olsB* mutant (**Figure S4F**). Colistin is a cyclic peptide that directly engages with the negative membrane charge, while the hydrophobic tail forms pores, leading to bactericidal activity. Colistin is a last resort antibiotic used against multidrug resistant Gram-negative bacterial infections. Our studies indicate increased colistin susceptibility in the *olsB*-deficient mutant relative to the wild-type strain (**Figure 4A and Figure S5A**), suggesting the antibiotic could be more effective against *A. baumannii* when aminolipid biosynthesis is inhibited. Under high phosphate conditions, the membrane lipid composition remains typical of standard bacterial membranes, resulting in higher relative colistin susceptibility (Figure 4C). However, in phosphate limited conditions, lipid turnover increases, leading to the replacement of phospholipids with non-phosphorylated aminolipids, which probably contributes to enhanced colistin tolerance. Overexpression of *olsB* in wild-type *A. baumannii* was sufficient to produce aminolipids during cultivation in excess phosphate (**Figure 4B**). Analysis demonstrated that *olsB* induction led to increased colistin tolerance (**Figure 4C and Figure S5B**). Therefore, OlsB-dependent aminolipid biosynthesis in *A. baumannii* promotes colistin tolerance, a last-line antibiotic for combating multidrug-resistant Gram-negative bacteria.

**Figure 4:**
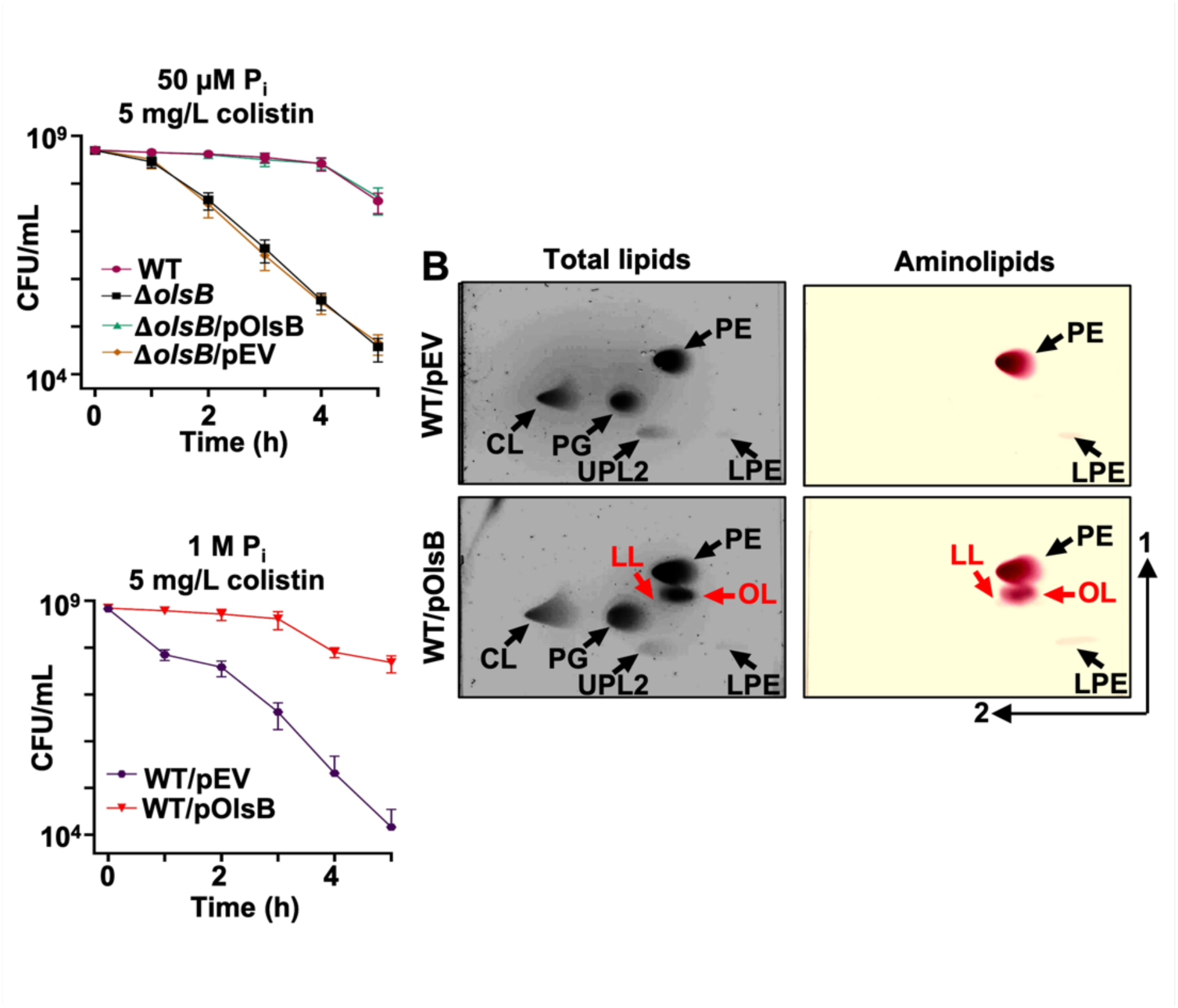
Aminolipids promote colistin tolerance in *A. baumannii*. **A.** Colistin-dependent killing in wild type and Δ*olsB* mutant strains under phosphate limitation (*n* = 3). Error bars indicate standard deviation. Wild type (WT) strains carrying pOlsB or empty vector were subjected to 5 mg/L colistin exposure over time. CFU/mL were calculated every 0.5 h. **B.** Total lipids were extracted using the Bligh and Dyer method and separated using 2D thin-layer chromatography. Lipids were stained with sulfuric acid (left). Aminolipids were stained using ninhydrin (right). Specific lipids are labelled: PE, phosphatidylethanolamine; PG, phosphatidylglycerol; CL, cardiolipin; LPE, lyso-phosphatidylethanolamine; LL, lysine lipid; OL, ornithine lipid; U3, unknown lipid 3. OL and LL are labelled in red. **C.** Colistin-dependent kill curves (left) and growth rate analysis (right) in wild type expressing empty vector (pEV) or pOlsB under high phosphate conditions (*n* = 3). Error bars indicate standard deviation.

### PhoR regulates aminolipid biosynthesis in *A. baumannii*

Previous work established that aminolipid production generally occurs in response to phosphate limitation (6,14,15,36,37). Consequently, we expected to find a Pho box upstream of the putative *olsB* gene. Using the *E. coli* pho box consensus sequence (38), a putative Pho box was identified that precedes the *olsB* gene in diverse *Acinetobacter* isolates, including ATCC 17978 (**Figure 5A**), but not preceding the *olsA* gene. The identified Pho box shows significant sequence similarity to the *E. coli* consensus (CT*GTCAT*NNNNC*TGTCAT*), with conserved nucleotides present in multiple *Acinetobacter* strains, including *A. baumannii* strains ATCC 17978, 19606, AB5075, AYE, and *A. baylyi*. These results are consistent with our transcriptomics analysis, which also suggested *olsA* gene expression is not responsive to phosphate concentrations.

**Figure 5:**
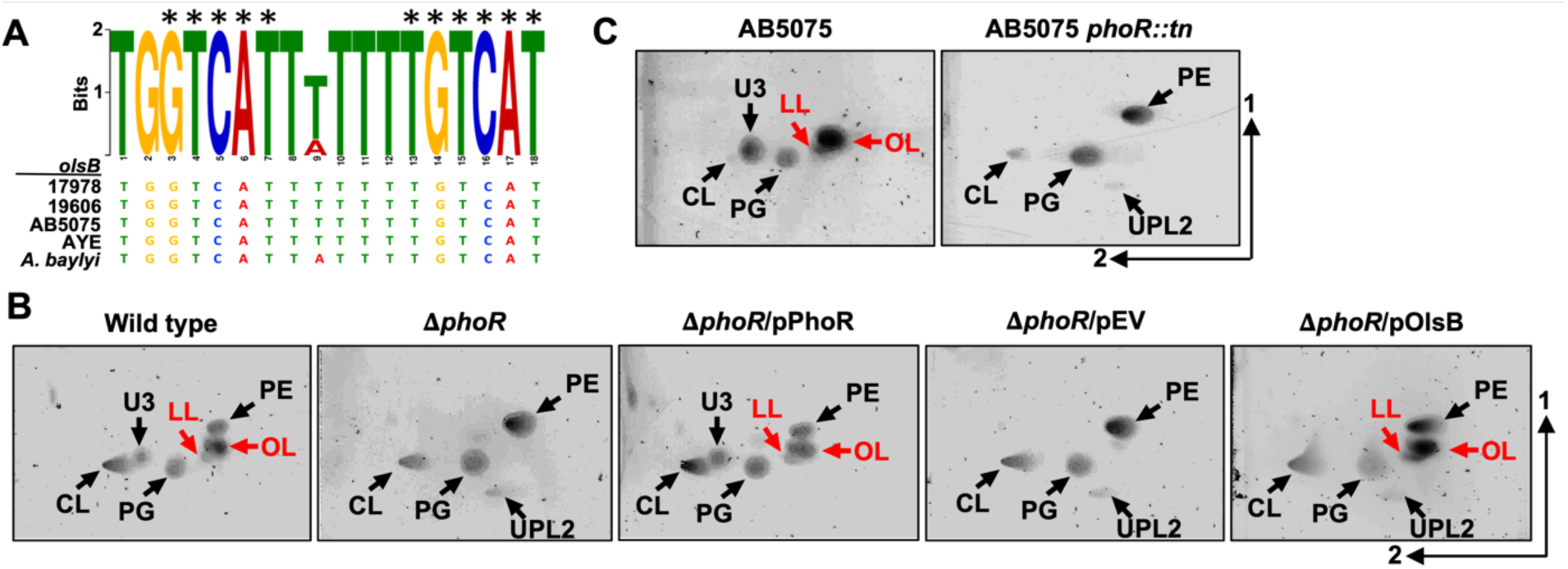
PhoR regulates *olsB* gene expression and aminolipid formation in *A. baumannii* strains. **A.** Predicted Pho box sequence alignments from *A. baumannii* strain 17978, 19606, AB5075, AYE, and *A. baylyi olsB* promoters. Black asterisks represent conserved nucleotides in the *E. coli* Pho box consensus (CT*GTCAT*NNNNC*TGTCAT*). **B.** 2D thin-layer chromatography lipid analysis from *A. baumannii* strain ATCC 17978 wild type and mutants grown in minimal media supplemented with 50 µM (limiting) phosphate. **C.** 2D thin-layer chromatography lipid analysis from *A. baumannii* strain AB5075 wild type and the *phoR* Tn*26* mutant in minimal media supplemented with 50 µM (limiting) phosphate. Lipids were stained with sulfuric acid. OL and LL aminolipids are labelled in red.

Phosphate sensing and diverse responses are regulated by the two-component system, PhoB/PhoR, in many bacteria (39). When phosphate levels decrease, the sensor kinase, PhoR, becomes activated, leading to autophosphorylation. Subsequent phosphotransfer to the cognate response regulator, PhoB results in a conformational change and DNA binding (40) at target promoters to induce gene expression.

These findings imply that the PhoB/PhoR two-component system regulates OL and LL biosynthesis in *A. baumannii* through *olsB* expression. To validate this hypothesis, we generated Δ*phoR* (*A1S_3376*) in *A. baumannii* ATCC 17978 by deleting codons 66 to 409. Lipid analysis after growth limiting phosphate, showed that the Δ*phoR* mutant accumulates glycerophospholipids PE, PG, CL, and UPL2, while failing to synthesize OL, LL, and U3, unlike the wild-type strain (**Figure 5B**). Complementation of the *phoR*-deficient mutant with the PhoR allele restored OL and LL formation, while aminolipids were absent in the mutant carrying the empty vector. Expression of *olsB* from a non-native promotor in the Δ*phoR* mutant restored OL and LL biosynthesis. Additionally, we examined the lipid patterns of the *phoR* (*ABUW_0105*)-transposon mutant of *A. baumannii* AB5075 grown under phosphate limitation (**Figure 5C**). Lipidomic analysis revealed that, unlike the wild-type strain, the *phoR::tn* mutant predominantly accumulated phospholipids and was unable to produce OL or LLs. Together, these findings suggest that *olsB* expression in *A. baumannii* is mediated by the *phoR* regulatory gene.

### Other Gram-negative ESKAPE pathogens form aminolipids under conditions of phosphate depletion

In addition to *P. aeruginosa* (14), *A. baumannii* is the second Gram-negative ESKAPE bacterium where the OL biosynthesis has been described using a OlsBA-dependent mechanisms. Uniquely, *A. baumannii* also produces LL via the same pathway. These findings prompted us to investigate if other Gram-negative ESKAPE pathogens, such as *Klebsiella* or *Enterobacter*, are also capable of aminolipid biosynthesis under phosphate limitation. TLC analysis of total lipids from *K. pneumoniae*, *E. cloacae* or *P. aeruginosa* after growth in minimal medium supplemented with excess phosphates showed production of the canonical membrane phospholipids PE, PG, and CL (**Figure 6A**). However, when cultivating *P. aeruginosa* in phosphate limitation (50 μM), a significant decrease in relative phospholipid levels was observed, and the synthesis of five unknown compounds and OL was induced. *K. pneumoniae* and *E. cloacae* cultivated in low phosphate concentrations (50 μM), also showed decreased levels of phospholipid production and accumulation of an unknown lipid. The unknown lipid exhibited a migration pattern and ninhydrin staining on TLC like OL, suggesting that aminolipid biosynthesis may be a conserved response to phosphate limiting growth conditions that could promote tolerance to antibiotics in Gram-negative ESKAPE pathogens.

**Figure 6:**
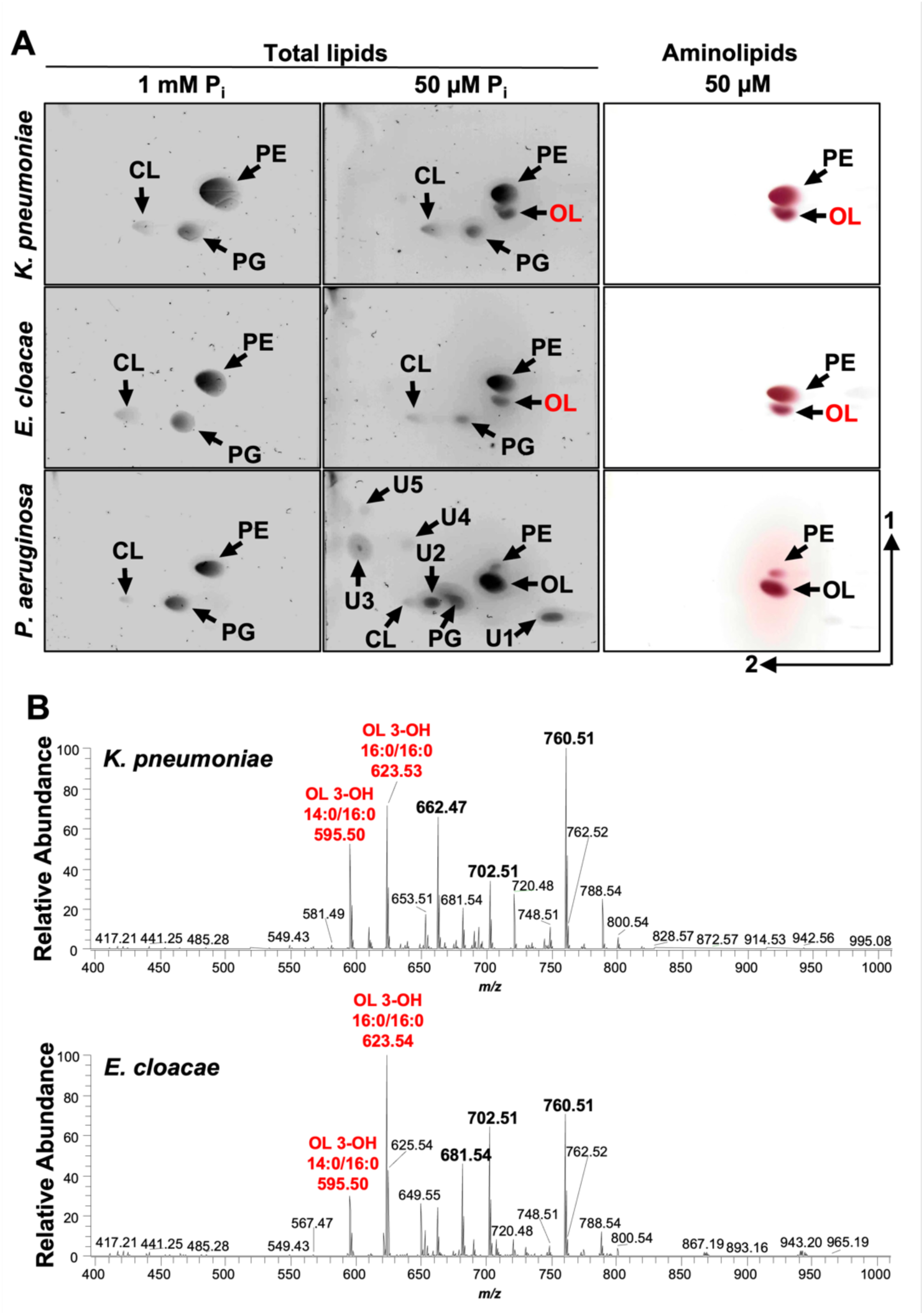
Aminolipid biosynthesis is conserved in Gram-negative ESKAPE pathogens. 2D thin-layer chromatography of lipids extracted from wild type *K. pneumoniae* strain KPNIH1, *E. cloacae* strain ATCC 13047, and *P. aeruginosa* strain PAO1 grown in excess (1 mM) or limiting (50 µM) phosphate conditions. Totals lipids were stained with sulfuric acid (left). Aminolipids were stained using ninhydrin (right). Specific lipids are labelled: PE, phosphatidylethanolamine; PG, phosphatidylglycerol; CL, cardiolipin; OL, ornithine lipid; U1, U2, U3, U4, and U5; unknown compounds 1, 2, 3, 4, and 5. **B.** Representative MS1 spectra from shotgun analysis of animolipids isolated from *K. pneumoniae* (*n* = 3) and *E. cloacae* (*n* = 3). The five most enriched lipids are bolded, including two ornithine lipids (OL), labelled in red.

To confirm the putative OL identity, total lipids were extracted using the Bligh and Dyer method after cells were grown in 50 μM phosphate. 1-D TLC (**Figure S6**) on total lipids stained with ninhydrin showed two prevalent lipids (lane 1). The band corresponding to OLs was scraped from the plate and extracted. Extracted lipids from three replicates were run on 1-D TLC showing enrichment of the putative OL relative to PE. Negative-ionization mode MS1 shotgun analysis on the purified lipids showed five enriched lipids in each strain (**Figure 6B**). MS/MS analysis of the lipids present in *K. pneumoniae* sample revealed two ornithine lipids (*m/z* 595.50 and 623.53), two PE lipids (*m/z* 662.47 and 702.51), and a *m/z* consistent with phosphatidylthreonine (*m/z* 760.51) (**Table S4**). Spectra from the *E. cloacae* lipid analysis included two ornithine lipids (*m/z* 623.54 and 595.50), PE (*m/z* 702.51), a *m/z* consistent with phosphatidylthreonine (*m/z* 760.51), and an unknown lipid that may be a modified OL (*m/z* 681.54) (**Table S5**). Together the migration of the lipids via 2-D TLC and the MS analysis provide strong complementary evidence that growth in limiting phosphate results in alternative synthesis of OLs.

## Discussion

Bacteria have evolved various regulatory mechanisms to sense and adapt to environmental stress. Here, we show that under phosphate-limiting conditions, *A. baumannii* produces ornithine and lysine lipids through regulated expression of *olsB*. The aminolipids alter the lipid bilayer composition to promote tolerance to colistin, an antimicrobial peptide used to treat Gram-negative bacterial infections. Specifically, aminolipids could reduce the electrostatic potential for cationic antimicrobial peptides such as colistin to target the cell, emphasizing changes in bilayer charge as an adaptive mechanism for *A. baumannii* in challenging environments. Although aminolipids are not directly exposed on the membrane surface, their accumulation in the inner leaflet of the outer membrane may alter the overall membrane dynamics, such as lipid packing and membrane fluidity, contributing to enhanced membrane stability and colistin tolerance. Broadly, cell membrane lipid modifications are vital for bacterial survival upon exposure to antimicrobial cationic peptides. For example, *V. cholerae* modifies lipid A with glycine or diglycine residues to resist cationic antimicrobial peptides (41). Similarly, other Gram-negative bacteria modify their lipid A structures to resist polymyxins. For instance, *P. aeruginosa* and *A. baumannii* can add positively charged amino groups, such as ethanolamine to lipid A, reducing to overall negative charge and decreasing the binding affinity of polymyxin (20). *K. pneumoniae* and *Enterobacter* species have been shown to add 4-amino-4-deoxy-L-arabinose (L-Ara4N) to lipid A, further neutralizing the negative charge and conferring resistance (21). Similar strategies are employed by various bacterial species, such as *P. aeruginosa*, *Rhizobium tropici*, *Staphylococcus aureus*, *Mycobacterium tuberculosis*, and *Bacillus subtilis*, which modify phospholipids like PG by adding amino acids like lysine or alanine, thereby conferring polymyxins resistance (36,37,42–44).

Membrane lipid remodeling during growth under phosphate limitation is a conserved strategy across bacteria, involving the substitution of phospholipids with phosphorus-free lipids alternatives like aminolipids (45). Interestingly, other bacteria, such as *P. aeruginosa*, also modify their lipid composition in response to phosphate limitation (14). Under these conditions, *P. aeruginosa* incorporates OL, replacing phospholipids in the membrane. This remodeling likely ensures that the membrane remains functional and stable despite the phosphate scarcity, highlighting a similar strategy employed by various bacterial species to adapt to nutrient stress. These findings further support the idea the replacement of phospholipids with amino acid-derived lipids, such as ornithine, represents an adaptive mechanism to phosphate limitation, suggesting that this process may be widely conserved across Gram-negative bacteria. While the synthesis of these lipids is not directly induced by the presence of polymyxins, tolerance to polymyxins is an indirect consequence of lipid membrane remodeling, leading to significant changes in membrane chemical properties. The alteration could reduce the net negative charge of the membrane, thereby decreasing CAMP susceptibility.

The *olsB* gene is required for OL biosynthesis, but unlike other OlsB-dependent pathways, it also induces LL formation in *A. baumannii*, suggesting metabolic plasticity within the species. Comparing our findings with observations in other bacteria, such as the soil bacterium *Rhodobacter sphaeroides*, which not only produces OLs but also synthesizes glutamine lipids (GLs) (46), further highlights the metabolic diversity in aminolipid biosynthesis. Additionally, recent research has shown that marine bacteria *Ruegeria pomeroyi* encode two *olsB* paralogs, one responsible for forming OLs and the other for GLs (30). Although these bacteria inhabit vastly different environments, the similarities suggest that the ability to use multiple substrates for aminolipid synthesis may be a common survival strategy in variable environmental conditions.

The transcriptomic analysis uncovered insights into different expression patterns of the *olsB* and *olsA* genes in response to phosphate limitation. Specifically, overexpression of the *olsB* gene was observed during phosphate limited growth, indicating a regulatory mechanism. However, *olsA* expression remained unchanged. The predicted Pho box in the promoter region of *olsB* suggested that the PhoB/PhoR two-component system regulates *olsB* under phosphate limitation conditions. The absence of a Pho box-like motif in the promotor region of *olsA*, in contrast to *olsB*, suggests that *olsA* is regulated by factors distinct from those controlling *olsB* under phosphate limiting conditions. This was confirmed by RNA sequencing (**Figure 3A**) and RT-PCR analysis (**Figure S4A**), which both showed that *olsA* expression remains unchanged in response to phosphate limitation, indicating that its expression is likely controlled by other regulatory mechanisms or constitutive. Consequently, *olsA* is not constitutively expressed under these conditions, but rather its regulation appears to be independent of phosphate availability. The absence of a similar motif in the promoter region of *olsA* implies the influence of other regulatory factors on its expression, independent of phosphate availability. Furthermore, *olsB* overexpression with non-native promoter in *A. baumannii* during cultivation in excess phosphate resulted in aminolipid formation, suggesting OlsA-dependent activity occurs after OlsB has formed a lyso-aminolipid. The differential regulatory events may also suggest that OlsA utilizes other substrates. For instance, in *Rhodobacter capsulatus*, OlsA functions as a bifunctional enzyme, active in both OL and phosphatidic acid biosynthesis (47). Additionally, we confirmed the role of the *phoR* gene as a key regulator in aminolipid biosynthesis in *A. baumannii*, highlighting its fine-tuned regulation in response to environmental stressors such as phosphate limitation. Interestingly, this regulatory mechanism shares similarities with species like *Sinorhizobium meliloti* or *V. cholerae*, where PhoB/PhoR also regulates OLs synthesis under phosphate limitation conditions (6,15), underscoring the evolutionary conservation of these adaptive mechanisms across different bacterial taxa.

Finally, the presence of *olsB* and *olsA* orthologues in *A. baumannii* raises questions about the diversity of aminolipid biosynthesis pathways among pathogens. Interestingly, bacteria such as *Klebsiella* and *Enterobacter* lack these orthologs, suggesting distinct biosynthesis pathways compared to *A. baumannii*. This absence prompts further investigation into alternative pathways or genes involved in aminolipid synthesis and their implications for antibiotics resistance and environmental adaptation.

### Conclusion

The study highlights the impact of phosphate limitation on lipid membrane composition in *A. baumannii*, resulting in the synthesis of OLs and LLs through regulated *olsB* gene expression. Aminolipids can promote tolerance to colistin, an important last-line antimicrobial. These results also underscore the role for *phoR* gene in regulating aminolipid synthesis in *A. baumannii* and show that other Gram-negative ESKAPE pathogens produce aminolipids under phosphate-depleted conditions. Aminolipid biosynthesis is a common adaptive response to phosphate limitation, which could promote pathogen survival both in hospital environment and within the host.

## Materials and methods

### Bacterial strains and growth

All strains and plasmids used in this study are listed in **Table S6** in the supplemental material. *E. coli*, *Acinetobacter* strains, *K. pneumoniae*, *E. cloacae* or *P. aeruginosa* were initially cultured from frozen stocks on Luria-Bertani (LB Miller) agar at 37°C. Isolated colonies were used to inoculate LB Miller broth or minimal medium (Tris minimal succinate [TMS]) (31); supplemented with different phosphate concentrations at 37°C. The minimal medium included: Na-succinate 20 mM, NaCl 200 mg/mL, NH_4_Cl 450 mg/mL, CaCl_2_ 200 mg/mL, KCl 200 mg/mL, MgCl_2_ 450 mg/mL, FeCl_2_ 10 mg/L, and MnCl_2_ 10 mg/L, with 10 mM 4-(2-hydroxyethyl)-1-piperazineethanesulfonic acid (HEPES) buffer used at pH 7.2. Na_2_HPO_4_ was then added to achieve a final concentration of 1 mM (excess) or 50 μM (limiting). All components were dissolved in deionized water and sterilized by filtration trough 0.22 μm pore-size filters and dissolved deionized H_2_O. To generate growth curves, overnight cultures of *A. baumannii* strains were diluted to an OD_600_ of ∼0.05 in 5 mL of medium, then incubated in glass test tubes at 37°C for 24 hours. Growth curve data were analyzed and plotted using GraphPad Prism software.

*A. baumannii* strain ATCC 17978 has been shown to be a mixture of two variants (48). Colony PCR using primers specific to the cardiolipin synthase gene (*clsC2*) resulted in a PCR product, indicating this study was specific to only the *A. baumannii* 17978 UN variant. *A. baumannii* strains AB5075 and ATCC 19606 have been shown to produce colonies with two opacity phenotypes that are phase variable (49). Microscopic examination showed a majority of opaque forms in strains AB5075 and ATCC 19606 with only 2-3 translucent colonies out of > 200 colonies found on each plate (*n* = 3 for each strain). The strain ATCC 19606 Δ*olsB* and strain AB5075 *olsB::tn* and *olsA::tn* mutants were also examined, but only included opaque variants.

### Construction of mutant and complementation *A. baumannii* strains

The primers utilized in this study are listed in **Table S7** in the supplementary material. Genetic mutants of *A. baumannii* were generated following established protocols (50–53). Briefly, *A. baumannii* carrying the pMMB67EH^TetR^ plasmid containing the REC*_Ab_* coding sequences was diluted from an overnight culture into LB Miller broth containing 10µg/mL of tetracycline at an OD_600_ of ∼0.05 and incubated for 45 minutes. REC*_Ab_* expression was induced by adding 2 mM IPTG, and cells were cultured at 37°C until they reached mid-log growth phase (OD_600_ of ∼0.4). After washing the cells three times in ice-cold 10% glycerol, 10^10^ cells were electroporated in a 2-mm cuvette at 1.8 mV with 5 µg of a recombineering linear PCR product. Subsequently, the cells were cultured for 4 hours in 4 mL of LB Miller broth with 2 mM IPTG and then plated on LB agar supplemented with 20 µg/mL of kanamycin. Mutations were validated using PCR.

To cure isolated mutants of the pMMB67EH^TetR^::REC*_Ab_* plasmid after mutant isolation, strains were streaked for isolated colonies on LB Miller agar supplemented with 2 mM NiCl_2_ to select cells that had lost the tetracycline cassette (51–54). Cured insertion mutants were then electroporated with pMMB67EH^TetR^ carrying the FLP recombinase. Cells were recovered for 1 hour in 5 mL of LB Miller broth and plated on LB agar containing 10µg/mL of tetracycline and 2 mM IPTG to induce expression of the FLP recombinase. Excision of the kanamycin cassette was confirmed by PCR.

To complement the *A. baumannii* ATCC 17978 mutants, the coding sequence from *A1S_0889* (*olsB*) was cloned into the KpnI/SalI sites, while the coding sequence from *A1S_3376* (*phoR*) was cloned into the KpnI/BamHI sites in pMMB67EH^KanR^. Plasmids were expressed in the respective mutants, and all strains were grown in 30 µg/mL of kanamycin and 1 mM IPTG for expression.

### Analysis of total lipids and aminolipids

Overnight cultures were used to inoculate 10 mL minimal medium 1 mM phosphate, or 20 mL minimal medium 50 µM phosphate to achieve an OD_600_ of ∼0.05, and then incubated for 24 hours at 37 °C. After the incubation period, cells were collected by centrifugation. Lipids were extracted using the Bligh and Dyer (1959) method (55). The chloroform phase was separated into the individual components on high-performance TLC silica gel plates. For one-dimensional TLC analysis, the plates were developed with chloroform-methanol-water (130:50:8 v/v) mixture. For two-dimensional TLC analysis, a chloroform-methanol-water (140:60:10 v/v) mixture was used in the first dimension and chloroform-methanol-acetic acid (130:50:20 v/v) mixture was used in the second dimension. Lipids on TLC were visualized by treating the plates with 10% sulfuric acid in ethanol at 150°C (total lipids) or 0.2% ninhydrin in acetone at 100°C (aminolipids). Lipid quantifications were done using ImageJ software to detect spots above background. Each experiment was independently replicated three times, one representative data set was reported in the quantification, and one representative image was included..

### Analysis of ^32^P-labeled phospholipids

Overnight cultures were diluted to an OD_600_ of ∼0.05 in 10 mL LB Miller broth supplemented with 5 µCi/mL ^32^P ortho-phosphoric acid (PerkinElmer) and grown until reaching an OD_600_ of ∼0.6. After harvesting the cells, lipid extraction was performed using Bligh and Dyer method (55), followed by analysis using TLC as previously described. Subsequently, the TLC plates were dried, exposed to a phosphorimaging screen, and scanned using an Amersham Typhoon laser scanner.

### Shotgun Mass Spectrometry

Dried lipid content from Bligh and Dyer extracted TLC bands of *E. cloacae* WT and *K. pneumoniae* WT biological replicates were solubilized in chloroform and resuspended in methanol: isopropanol (1:1) at a concentration of 200 µM. Shotgun analysis was performed in negative-ionization mode on a Thermo Scientific Orbitrap Fusion Lumos Tribrid mass spectrometer. Nanoelectrospray ionization was performed using Pd/Au-coated pulled-tip glass capillaries (20 nm i.d.) at a spray voltage of - 1000 V and a capillary temperature of 275°C. Spectra were acquired at a resolution of 50,000 at *m/z* 200, an AGC target between 5E4 and 1E5, and an RF lens of 60%. All MS1 and MS/MS spectra were an average of 50 scans. The top 5 most abundant species were subjected to higher energy collisional dissociation (HCD) for identification. MS/MS spectra were obtained with an isolation width of 1 *m/z* and a normalized collision energy (NCE) ranging from 22-26%.

### Liquid chromatography

Lipids isolated from *Acinetobacter baumannii* were separated using reversed-phase liquid chromatography (LC) on an Acquity UPLC CSH C18 column (pore size 130 Å, 1.7 μm particle size, 2.1 mm × 100 mm, Waters) integrated with a Dionex Ultimate 3000 UHPLC system (Thermo Fisher Scientific). Mobile phase A and B were comprised of methanol: water: acetonitrile (3:4:3) and isopropanol: water: acetonitrile (90:2.5:7.5), respectively, each containing 10 mM ammonium formate. Dried lipid content from TLC bands was extracted via a Bligh and Dyer protocol and resuspended in 50:50 mobile phase (A: B) at a concentration of 100 ng/µL. A 9 µL injection volume was used, and the column compartment was maintained at a temperature of 50 °C. Aminolipids were separated at a flow rate of 0.275 mL/min with the following gradient: hold at 10% B (0-1 min), 10-45% B (1-5 min), 45-70% B (5-23 min), 70-95% B (23-24 min), hold at 95% B (24-29 min), 95-10% B (29-29.5), and hold at 10% B (29.5-35 min).

### Untargeted LC-MS with HCD and Targeted LC-MS with UVPD Experiments

An untargeted negative-ionization mode LC-MS with HCD method was utilized to screen *Acinetobacter baumannii* lipid extracts on a Thermo Scientific Orbitrap Fusion Lumos Tribrid mass spectrometer via heated electrospray ionization. Various source parameters included a spray voltage of -3800 V, sheath gas setting of 5, aux gas setting of 10, and capillary temperature of 300 °C. MS1 data was collected at a resolution of 30,000 at *m/z* 200 in the Orbitrap analyzer with a scan range of *m/z* 500-1000, RF lens of 80%, and an AGC target of 5E5. A data-dependent acquisition method was used where the top 5 most abundant ions above an intensity threshold of 5E4 were selected for MS/MS analysis with HCD. Species were isolated using a 1 *m/z* window and subjected to a collision energy corresponding to NCE of 22%. MS/MS spectra were acquired at a resolution of 30,000 at *m/z* 200 in the Orbitrap analyzer, AGC target of 1E5, maximum injection time of 250 ms, and 2 microscans/scan.

A targeted positive-ionization mode LC-MS/MS method was performed on a Thermo Scientific Orbitrap Eclipse Tribrid mass spectrometer equipped with an ArF excimer laser (Coherent, Inc.) for 193 nm ultraviolet photodissociation (UVPD). The heated electrospray ionization source was operated at + 3800 V, and the source parameters described in the untargeted experiments were implemented. MS/MS scans with UVPD were acquired in a data-dependent manner, with 5 scans collected between each MS1 master scan, and with an intensity threshold set at 5E4. A targeted mass filter was employed that included the *m/z* values and start/end retention times for unsaturated aminolipids identified from the negative-ionization mode LC-MS with HCD run (with a 15-ppm error tolerance). Aminolipids were isolated with a 1 *m/z* window and activated with 4 laser pulses at 2 mJ/pulse for double bond localization. MS/MS spectra were acquired at a resolution of 30,000 at *m/z* 200 in the Orbitrap analyzer, q-value of 0.1, AGC target of 5E5, maximum injection time of 500 ms, and 5 microscans/scan. Data was manually interpreted using Thermo Xcaliber Qual Browser and ChemDraw. Precursor and fragment ions were assigned within 10 and 15 ppm mass error, respectively.

### RNA sequencing

Transcriptome sequencing analysis was performed as described previously, with modification (53). Briefly, total RNA was extracted from *A. baumannii* ATCC 17978 cultures grown in minimal medium supplemented with either excess (1 mM) or limiting (50 μM) phosphate at OD_600_ of ∼0.5 in triplicate, utilizing the Direct-Zol RNA miniprep kit (Zymo Research). Genomic DNA contamination was eliminated using the Turbo DNA-free DNA removal kit (Invitrogen). DNAase-treated RNA samples were then forwarded to SeqCenter for sequencing on the Illumina NextSeq 550 sequencing. Subsequently, the CLC Genomic Workbench software (Qiagen) was employed to map the obtained sequencing data to the *A. baumannii* ATCC 17978 genome annotations and determine the read per kilobase per million (RPKM) expression values and determine the weighted-proportions fold changes in expression values between excess or limitation phosphate conditions. Data were analyzed and plotted using GraphPad Prism software. Data Accession #: GSE276010.

### Quantitative RT-PCR

Relative-abundance quantitative PCR (qPCR) was performed as previously described In brief, total RNA was extracted from bacterial cultures using the Direct-Zol RNA Miniprep Kit (Zymo Research), followed by DNase digestion to remove any genomic DNA. The RNA was then reverse transcribed into cDNA using the Invitrogen SuperScript III Reverse Transcriptase (Thermo Fisher), with a final RNA concentration of 5 ng/µl. One microliter of the resulting cDNA was used as a template in a 10-µl quantitative RT-PCR (qRT-PCR) reaction, conducted with Power SYBR Green Master Mix (Applied Biosystems). qRT-PCR was carried out on a QuantStudio 3 Real-Time PCR System (Thermo Fisher Scientific). Relative expression levels were calculated using the ΔΔCt method (56), with normalization of gene targets to the *rpoA* signal.

### Microscopy and image analysis

Cells were grown as stated above and fixed with paraformaldehyde (PFA) and mounted on 1.5% agarose in 1X phosphate-buffered saline (PBS). Imaging was performed using a Nikon Eclipse Ti-2 wide-field epifluorescence microscope equipped with a Photometrics Prime 95B camera and a Plan Apo 100x, 1.45 numerical aperture objective lens. Images were captured using NIS Elements software. Image analysis was conducted using the microbeJ plugin of ImageJ software.

### LOS staining and analysis

All cultures were grown in test tubes containing 5 mL of minimal medium with either excess (1 mM) or limiting (50 μM) phosphate at 37°C in a shaker overnight. For the complementation strains, 30 µg/mL kanamycin and 1 mM IPTG were added. The OD_600_ of the overnight cultures was measured and normalized to OD_600_ of ∼1. The cells were then centrifuged at 15,000 rpm for 5 minutes. Each pellet was resuspended in 100 μl of 1X Sample Buffer (4X LDS Sample Buffer, 4% β-mercaptoethanol, and water) and boiled in water for 10 minutes. After cooling, proteinase K was added to each sample and mixed. The samples were then incubated in a 55°C water bath overnight. The following day, the samples were boiled in water for 5 minutes and SDS-PAGE was performed. The gel was then fixed and treated according to the protocol in the Pro-Q Emerald 300 Lipopolysaccharide Gel Stain Kit by Thermo Fisher Scientific (P20495).

### Colistin susceptibility assays

Overnight cultures were diluted to OD_600_ ∼0.150 in minimal medium supplemented with either excess (1 mM) or limiting (50 μM) phosphate containing 5 mg/L colistin. Each culture, comprising 15 mL, was incubated in 125 mL Erlenmeyer flasks at 37°C with agitation at 250 rpm. Survivors were analyzed at specific time points by serial dilution plating on LB agar.

## Supplementary Captions

**Figure S1: *Escherichia coli* (*Ec*) and *Acinetobacter baumannii (Ab)* lipid composition after growth in complex media. A.** Strains were grown in complex media (LB broth) in the presence of ^32^P-orthopohosphoric acid until mid-logarithmic growth phase. Cells were collected and total lipids were extracted using the Bligh and Dyer method and separated using 2-dimensional thin-layer chromatography. Labelled lipids include anionic cardiolipin (CL) and phosphatidylglycerol (PG), aminolipids phosphatidylethanolamine (PE) and lyso-PE (LPE), and unknown phospholipid 1 (UPL1). **B.** PG, PE, CL and LPE chemical structures. Head groups are red, the glycerol backbone is blue, and fatty acids are black.

**Figure S2: Effect of phosphate availability on growth, cell morphology, and LOS production. A.** Quantification of total lipid in TLC (A) grown in excess or limiting phosphate concentrations. Lipids are graphed as a percentage of the total. Significance testing conducted using Student *t* test with two-tailed distribution assuming equal variance. Lines indicate standard deviation. \**P* < 0.05; ns is not significant. **B.** *A. baumannii* ATCC 17978 was cultured in minimal medium containing either excess (1 mM) or limiting (50 µM) phosphate for 24 hours. **C.** Phase-contrast images of *A. baumannii* grown under these conditions, captured during the exponential phase. Scalebar is 10 µm. **D.** Cell length and area were quantified for each population (*n* ≥ 150) using ImageJ software. Each point represents an individual cell. The experiment was repeated twice, and one representative dataset was reported. Significance testing conducted using Student *t* test with two-tailed distribution assuming equal variance. \*\*\**P* < 0.001, \*\*\*\**P* < 0.0001. **E.** Proteinase K-treated whole-cell lysate from wild-type *A. baumannii* strains ATCC 17978, ATCC 19606, and AB5075, as well as from aminolipid-deficient mutants grown under excess (1mM) or limiting (50 µM) phosphate conditions. The LOS samples were separated using SDS-PAGE and stained using Pro-Q Emerald 300.

**Figure S3: Thin-layer chromatography of *Acinetobacter* aminolipids. A.** To produce *A. baumannii* lipid samples for MS analysis, total lipids were separated using thin-layer chromatography and scraped from the plate, extracted using the Bligh and Dyer method, and run alongside lipid controls. Extractions resulted in isolation of U1 & U2, U1, or U2. Lipids were stained with ninhydrin to visualize aminolipids. Specific lipids are labelled: PE, phosphatidylethanolamine; U1, unknown lipid 1; U2, unknown lipid 2. **B.** Cells were grown in minimal medium supplemented with excess (1mM) or limiting (50 µM) phosphate conditions from indicated *Acinetobacter* strains. Total lipids were extracted from cells grown in media with limiting phosphate concentrations. Total lipids were spotted on thin-layer chromatography and separated based on hydrophobicity. Plates were stained with ninhydrin to visualize aminolipids. Specific lipids are labelled: PE, phosphatidylethanolamine; OL, ornithine lipid; LL, lysine lipid. **C.** To produce *E. cloacae* and *K. pneumoniae* samples for MS analysis, total lipids were extracted from cells grown in media with limiting phosphate concentrations. Total lipids were separated using thin-layer chromatography and scraped from the plate, extracted using the Bligh and Dyer method, and run alongside lipid controls. Lipids were stained with ninhydrin to visualize aminolipids. Specific lipids are labelled: PE, phosphatidylethanolamine; OL; ornithine lipid (predicted).

**Figure S4: *olsB* and *olsA* are required for ornithine and lysine lipid biosynthesis in *A. baumannii*. A.** Relative-abundance quantitative PCR (qPCR) of genes after *A. baumannii* growth in excess or limiting phosphate concentrations (*n* = 3). Lines indicate standard deviation. Significance testing was conducted using Student *t* test with two-tailed distribution assuming equal variance. \**P* < 0.05; ns = not significant. **B.** 2D thin-layer chromatography lipid analysis in ATCC 19606 wild type and the Δ*olsB* mutant strain after growth in limiting (50 µM) phosphate concentrations. Lipids were stained with sulfuric acid. **C.** Analysis in AB5075 wild type and transposon (Tn*101*) mutant strains after growth in limiting (50 µM) phosphate concentrations. Specific lipids are labelled: PE, phosphatidylethanolamine; PG, phosphatidylglycerol; CL, cardiolipin; OL, ornithine lipid; LL, lysine lipid. OL and LL aminolipids are labelled in red. **D.** Phase-contrast images of *A. baumannii* strain ATCC 17978 grown under phosphate limiting conditions, captured during exponential phase growth. Scalebar is 10 µm. **E.** Cell length and area of strain ATCC 17978 were quantified for each population (*n* ≥ 150) using ImageJ software. Each point represents an individual cell. Significance testing was conducted using Student *t* test with two-tailed distribution assuming equal variance. ns = not significant. **F**. Optical density (OD_600_) measurements of *A. baumannii* strains grown in 50 µM phosphate over 24 h.

**Figure S5: Aminolipid formation promotes *A. baumannii* tolerance to colistin.** Growth (OD_600_) of *A. baumannii* ATCC 17978 strains was measured at 37 °C in minimal medium with limiting phosphate (**A**) or excess phosphate (**B**), and in the presence of colistin at concentrations of 1, 2.5, or 5 mg/L.

**Figure S6: Thin-layer chromatography of *E. cloacae* and *K. pneumoniae* aminolipids.** To produce *E. cloacae* and *K. pneumoniae* samples for MS analysis, total lipids were extracted from cells grown in media with limiting phosphate concentrations. Total lipids were separated using thin-layer chromatography and scraped from the plate, extracted using the Bligh and Dyer method, and run alongside lipid controls. Lipids were stained with ninhydrin to visualize aminolipids. Specific lipids are labelled: PE, phosphatidylethanolamine; OL; ornithine lipid (predicted).

**Table S1: Identity of aminolipids in unknown lipid 1 biological replicates.**

**Table S2: Identity of aminolipids in unknown lipid 2 (U2) biological replicates.**

**Table S3: Differentially regulated genes in excess and limiting phosphate**

**Table S4: Lipids present in the top five most abundant species from *K. pneumoniae* biological replicates**

**Table S5: Lipids present in the top five most abundant species from *E. cloacae* biological replicates**

**Table S6: Strains and plasmids used in this study.**

**Table S7: Primers used in this study.**

