## Supplemental figures 1-6 and tables 6 and 7 for "Alternative lipid synthesis in response to phosphate limitation promotes antibiotic tolerance in Gram-negative ESKAPE pathogens"

### Supplementary Figures

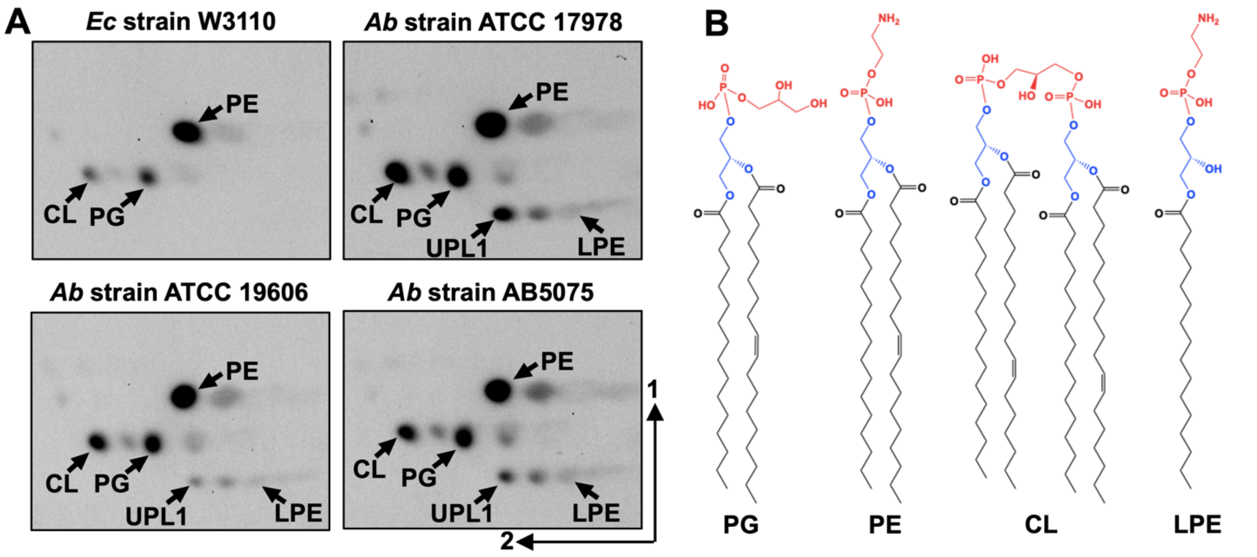

**Figure S1: *Escherichia coli* (*Ec*) and *Acinetobacter baumannii (Ab)* lipid composition after growth in complex media. A.** Strains were grown in complex media (LB broth) in the presence of $^{32}$P-orthopohosphoric acid until mid-logarithmic growth phase. Cells were collected and total lipids were extracted using the Bligh and Dyer method and separated using 2-dimensional thin-layer chromatography. Labelled lipids include anionic cardiolipin (CL) and phosphatidylglycerol (PG), aminolipids phosphatidylethanolamine (PE) and lyso-PE (LPE), and unknown phospholipid 1 (UPL1). **B.** PG, PE, CL and LPE chemical structures. Head groups are red, the glycerol backbone is blue, and fatty acids are black.

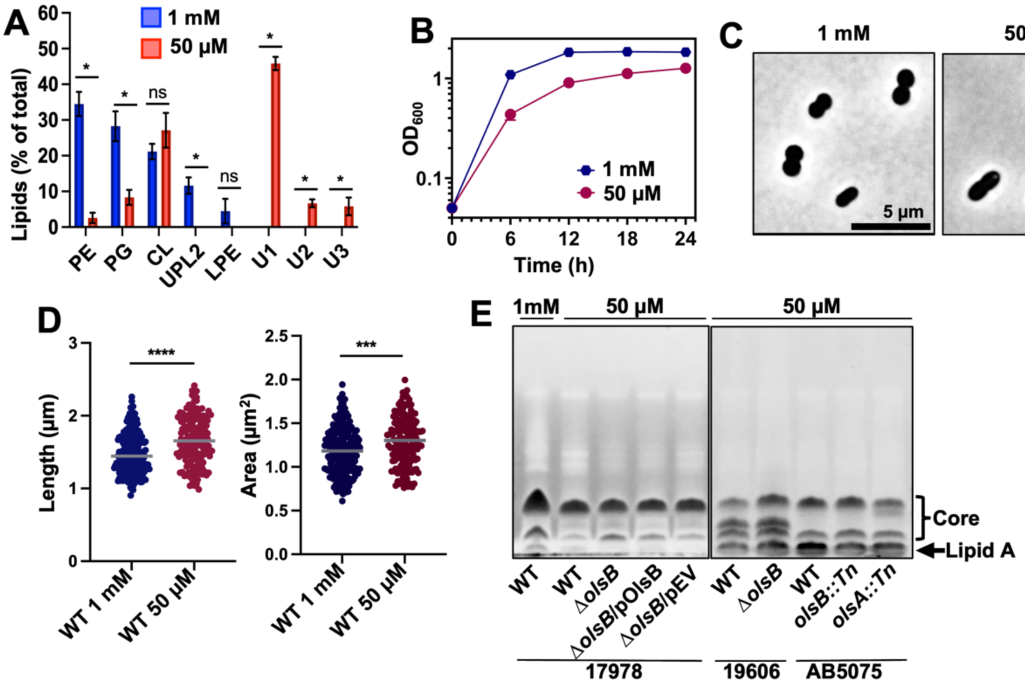

**Figure S2: Effect of phosphate availability on growth, cell morphology, and LOS production. A.** Quantification of total lipid in TLC (A) grown in excess or limiting phosphate concentrations. Lipids are graphed as a percentage of the total. Significance testing conducted using Student *t* test with two-tailed distribution assuming equal variance. Lines indicate standard deviation. *$P < 0.05$; ns is not significant. **B.** *A. baumannii* ATCC 17978 was cultured in minimal medium containing either excess (1 mM) or limiting (50 µM) phosphate for 24 hours. **C.** Phase-contrast images of *A. baumannii* grown under these conditions, captured during the exponential phase. Scalebar is 10 µm. **D.** Cell length and area were quantified for each population ($n \geq 150$) using ImageJ software. Each point represents an individual cell. The experiment was repeated twice, and one representative dataset was reported. Significance testing conducted using Student *t* test with two-tailed distribution assuming equal variance. ***$P < 0.001$, ****$P < 0.0001$. **E.** Proteinase K-treated whole-cell lysate from wild-type *A. baumannii* strains ATCC 17978, ATCC 19606, and AB5075, as well as from aminolipid-deficient mutants grown under excess (1mM) or limiting (50 µM) phosphate conditions. The LOS samples were separated using SDS-PAGE and stained using Pro-Q Emerald 300.

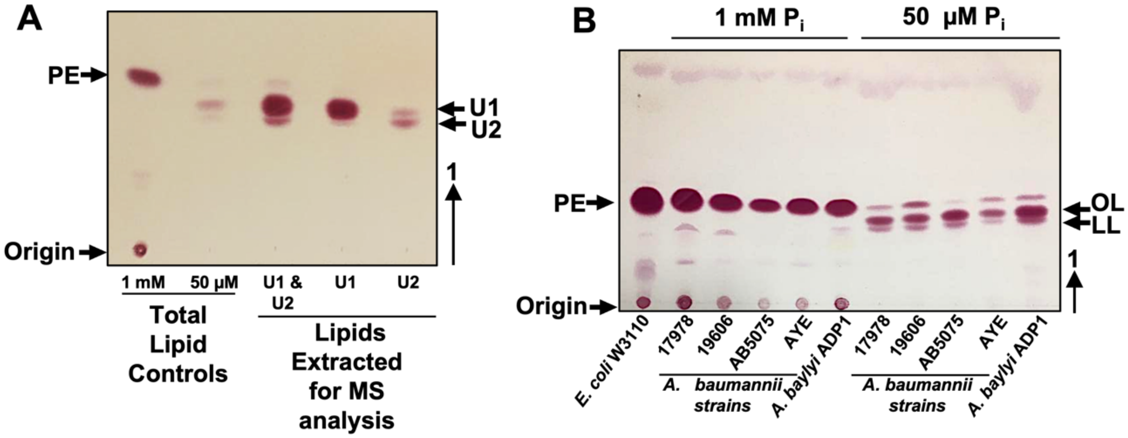

**Figure S3: Thin-layer chromatography of *Acinetobacter* aminolipids. A.** To produce *A. baumannii* lipid samples for MS analysis, total lipids were separated using thin-layer chromatography and scraped from the plate, extracted using the Bligh and Dyer method, and run alongside lipid controls. Extractions resulted in isolation of U1 & U2, U1, or U2. Lipids were stained with ninhydrin to visualize aminolipids. Specific lipids are labelled: PE, phosphatidylethanolamine; U1, unknown lipid 1; U2, unknown lipid 2. **B.** Cells were grown in minimal medium supplemented with excess (1mM) or limiting (50 µM) phosphate conditions from indicated *Acinetobacter* strains. Total lipids were extracted from cells grown in media with limiting phosphate concentrations. Total lipids were spotted on thin-layer chromatography and separated based on hydrophobicity. Plates were stained with ninhydrin to visualize aminolipids. Specific lipids are labelled: PE, phosphatidylethanolamine; OL, ornithine lipid; LL, lysine lipid. **C.** To produce *E. cloacae* and *K. pneumoniae* samples for MS analysis, total lipids were extracted from cells grown in media with limiting phosphate concentrations. Total lipids were separated using thin-layer chromatography and scraped from the plate, extracted using the Bligh and Dyer method, and run alongside lipid controls. Lipids were stained with ninhydrin to visualize aminolipids. Specific lipids are labelled: PE, phosphatidylethanolamine; OL; ornithine lipid (predicted).

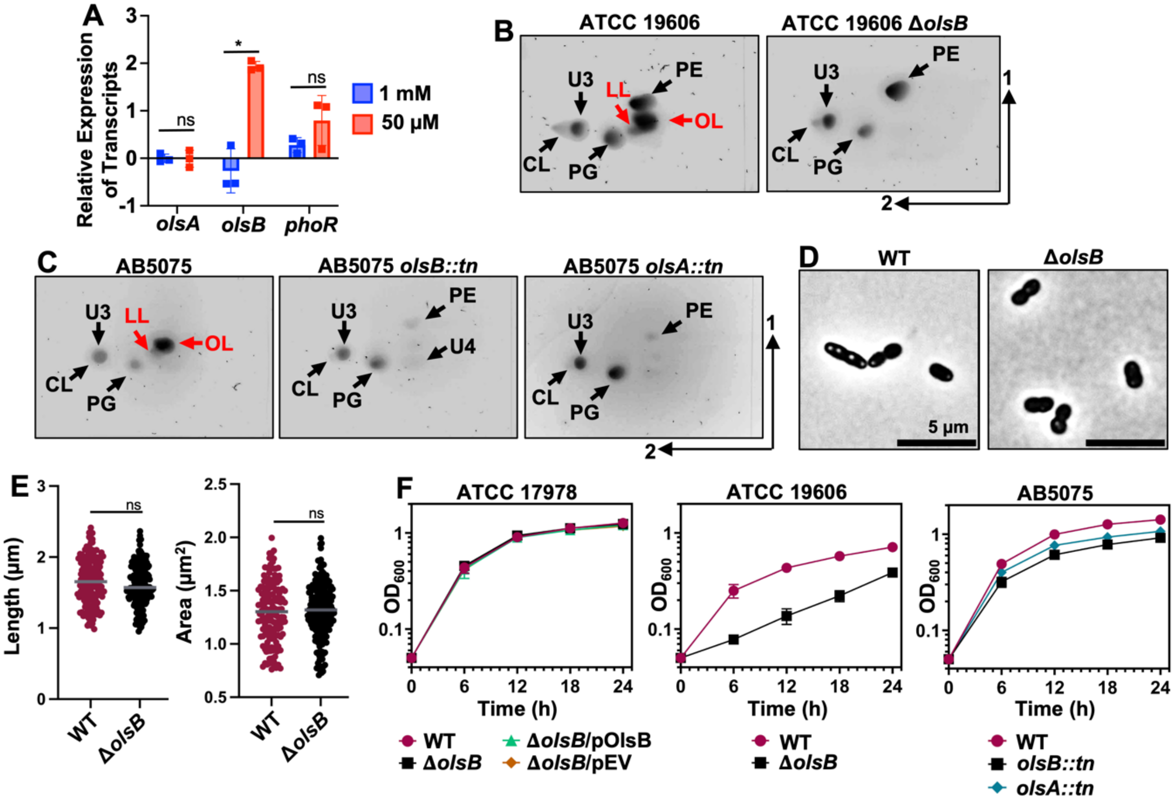

**Figure S4: *olsB* and *olsA* are required for ornithine and lysine lipid biosynthesis in *A. baumannii*.** **A.** Relative-abundance quantitative PCR (qPCR) of genes after *A. baumannii* growth in excess or limiting phosphate concentrations ($n$ = 3). Lines indicate standard deviation. Significance testing was conducted using Student $t$ test with two-tailed distribution assuming equal variance. *$P < 0.05$; ns = not significant. **B.** 2D thin-layer chromatography lipid analysis in ATCC 19606 wild type and the Δ*olsB* mutant strain after growth in limiting (50 µM) phosphate concentrations. Lipids were stained with sulfuric acid. **C.** Analysis in AB5075 wild type and transposon (Tn*101*) mutant strains after growth in limiting (50 µM) phosphate concentrations. Specific lipids are labelled: PE, phosphatidylethanolamine; PG, phosphatidylglycerol; CL, cardiolipin; OL, ornithine lipid; LL, lysine lipid. OL and LL aminolipids are labelled in red. **D.** Phase-contrast images of *A. baumannii* strain ATCC 17978 grown under phosphate limiting conditions, captured during exponential phase growth. Scalebar is 10 µm. **E.** Cell length and area of strain ATCC 17978 were quantified for each population ($n \geq 150$) using ImageJ software. Each point represents an individual cell. Significance testing was conducted using Student $t$ test with two-tailed distribution assuming equal variance. ns = not significant. **F.** Optical density ($OD_{600}$) measurements of *A. baumannii* strains grown in 50 µM phosphate over 24 h.

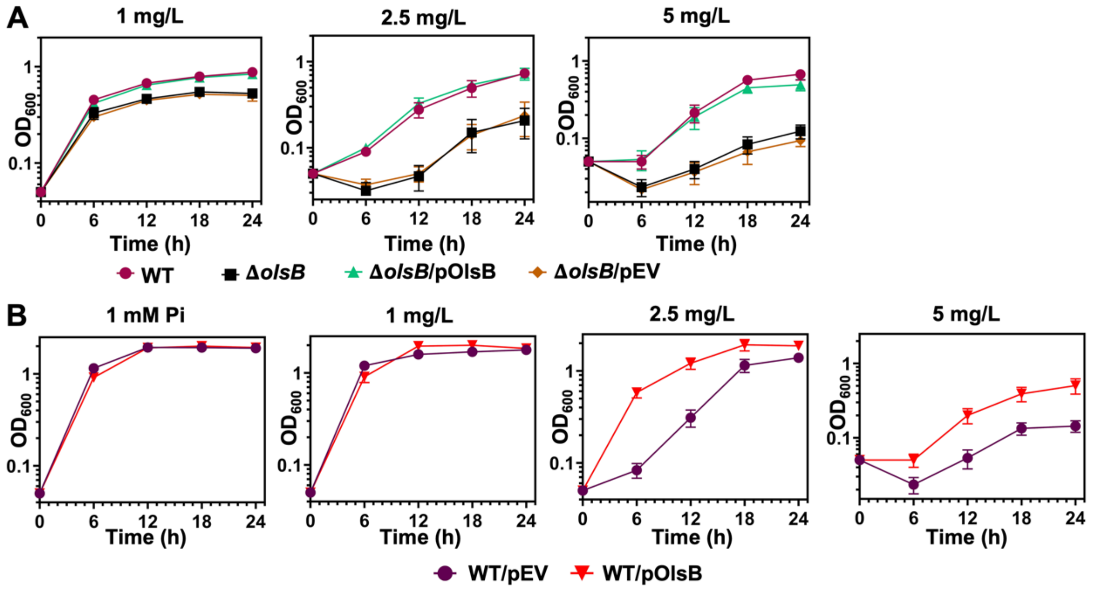

**Figure S5: Aminolipid formation promotes *A. baumannii* tolerance to colistin.** Growth ($OD_{600}$) of *A. baumannii* ATCC 17978 strains was measured at 37 °C in minimal medium with limiting phosphate (**A**) or excess phosphate (**B**), and in the presence of colistin at concentrations of 1, 2.5, or 5 mg/L.

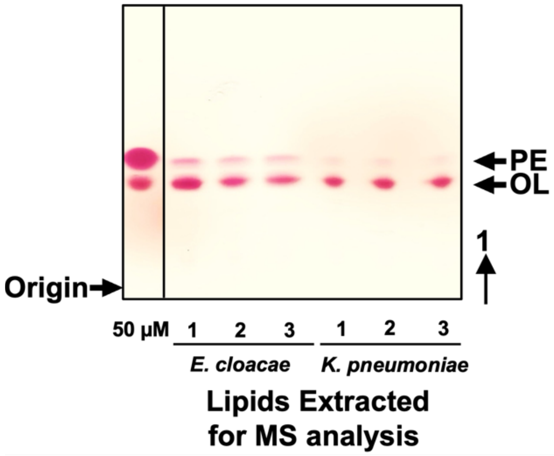

**Figure S6: Thin-layer chromatography of *E. cloacae* and *K. pneumoniae* aminolipids.** To produce *E. cloacae* and *K. pneumoniae* samples for MS analysis, total lipids were extracted from cells grown in media with limiting phosphate concentrations. Total lipids were separated using thin-layer chromatography and scraped from the plate, extracted using the Bligh and Dyer method, and run alongside lipid controls. Lipids were stained with ninhydrin to visualize aminolipids. Specific lipids are labelled: PE, phosphatidylethanolamine; OL; ornithine lipid (predicted).

**Supplementary Tables**

**Please See the Excel Spreadsheet for Tables S1-3**

**Table S6: Strains and plasmids used in this study.**

| Strain or Plasmid | Description | Reference or Source |
|---|---|---|
| **Strains** | | |
| *A. baumannii* ATCC 17978 | Wild type | (1) |
| *A. baumannii* ATCC 19606 | Wild type | (2) |
| *A. baumannii* AB5075 | Wild type | (3) |
| *A. baumannii* AYE | Wild type | (4) |
| *A. baylyi* ADP1 | Wild type | (5) |
| *P. aeruginosa* PAO1 | Wild type | (6) |
| *K. pneumoniae* KPNIH1 | Wild type | (7) |
| *E. cloacae* ATCC 13047 | Wild type | (8) |
| *A. baumannii* ATCC 17978 | *ΔolsB (A1S_0889)::deletion* | This study |
| *A. baumannii* ATCC 17978 | *ΔphoR (A1S_3376)::deletion* | This study |
| *A. baumannii* ATCC 19606 | *ΔolsB (HMPREF0010_01383)::deletion* | This study |
| *A. baumannii* AB5075 | *ΔolsB (ABUW_3039)::tn101* | (9) |
| *A. baumannii* AB5075 | *ΔolsA (ABUW_0502)::tn101* | (9) |
| *A. baumannii* AB5075 | *ΔphoR (ABUW_0105)::tn26* | (9) |
| *E. coli* W3110 | Wild type, F- *λ-*, *rph-1 IN (rrnD, rrnE)1* | (10) |
| *E. coli* DH5α | recA1, ϕ80 *lacZ*ΔM15, host for cloning | (11) |
| **Plasmids** | | |
| pAT03 | pMMB67EH with FLP recombinase, Tet$^{R}$ | (12) |
| pAT04 | pMMB67EH with REC$_{Ab}$ system, Tet$^{R}$ | (12) |
| pKD4 | Kan$^{R}$ | (13) |

| | | |
|---|---|---|
| pMMB67EHKn | pMMB67EH with the Kan$^{R}$ gene from pKD4 inserted into the PvuI site, Kan$^{R}$ | (14) |
| pROB01 | pMMB67EHKn carrying *olsB* (*A1S_0889*) | This study |
| pROB02 | pMMB67EHKn carrying *phoR* (*A1S_3376*) | This study |

**Table S7: Primers used in this study.**

| **Oligo Name** | **Sequence (5' to 3')** |
|---|---|
| **Deletion Primers** | |
| 17978 *olsB*Ab Kan FRT 5' | AGTGGGTGTCACTAGGAGCGTTCATTATGCTGGAAAAATTTAATCAATATCGCCAAACCTGGACTTTACCTTTAAATCGCCATAAGGCTAACAATCAAACACAATTCCGCTTTGAATGGGTTGATAGCGATTGTGTAGGCTGGAGCTGCTTCG |
| 17978 *olsB*Ab Kan FRT 3' | TTGTTCATTACAAACGAAGTGGCAATTTTATTCACTTCTAAAAATACAAAGTAATCGAGACAGTTAAATTCAGCATCAAAGAAAGCATCTTTAGATAATTTAGACTGCATACTCAAATACATTTGATATCCTCCTTAGTTCCTATTCCG |
| 17978 *olsB*Ab confirm 5' | GTGGTGTTGTGAGCGCACATATTG |
| 17978 *olsB*Ab confirm 3' | GCCTTGAGTCGCCTTACGAATATG |
| 17978 *phoR*Ab Kan FRT 5' | CGTTTGCTAAACAAGATTTACGACTTTTATTATTTTTCCTGATTATTGCAGGTTTAGTCGGTTTAGGAATTGGGTATTTCTGGAGCTGTATTTTTATTGCCTTTGTGGTGTTTTTTACACTTCAGAGCGATTGTGTAGGCTGGAGCTGCTTCG |
| 17978 *phoR*Ab Kan FRT 3' | ATGTTATAGAGTCTTTCTTTTGGAAAAACTGCGGTAAAGGTTGATCCTTCATTTTCTTTAGATTGCACATCTAAGTAGGCGCCGTGTTGCATGAGTACATGTTTTACAATCGCCAAGCCTAAACCATATCCTCCTTAGTTCCTATTCCG |
| 17978 *phoR*Ab 5' confirm | GGACCAACAGAATACCGTCTGCTTG |
| 17978 *phoR*Ab 3' confirm | GATGGTGGAGATCATCGTGATGCAC |
| **Complementation Primers** | |
| 17978 *olsB*Ab KpnI 5' | CGCGGTACCATGCTGGAAAAATTTAATCAATATCGC |
| 17978 *olsB*Ab SalI 3' | CGCGTCGACTTATCGCTGAGCCATTTTGTTC |
| 17978 *phoR*Ab SacI 5' | CGCGAGCTCATGTATGAACCCTACCCCGTCC |
| 17978 *phoR*Ab BamHI 3' | CGCGGATCCTTAAGTCATGTTATAGAGTCTTTCTTTTGG |
| **Transposon mutant primers** | |
| AB5075 *olsB::tn101* 5' | CGACACAAGAAGTTGACCGTTTAATCG |

| | |
|---|---|
| confirm | |
| AB5075 *olsB::tn101* 3' confirm | CAATTGGTGATTCAACTGCAGACTGG |
| AB5075 *olsA::tn101* 5' confirm | GCTCAAGAATATCTGCACCAGCTGAC |
| AB5075 *olsA::tn101* 3' confirm | GGATGAGCTGGCAACTAAAGCG |
| AB5075 *phoR::tn26* 5' confirm | GTCTGGATGCTGGTGCAGATGAC |
| AB5075 *phoR::tn26* 3' confirm | GGTGCCGACTGTACCGTTAATG |
